# Opposing sleep and feeding phenotypes across development in desert flies

**DOI:** 10.1101/2025.10.15.682659

**Authors:** Shuhao Li, Milan Szuperak, Ceazar Nave, Si Hao Tang, Jeffrey M. Donlea, Matthew S. Kayser

## Abstract

Sleep is essential for adaptation and survival across the lifespan, yet the ecological pressures shaping sleep ontogeny remain poorly understood. We investigated sleep across early developmental stages in *Drosophila mojavensis*, a stress-resilient, desert-adapted species. While adult *D. mojavensis* exhibit prolonged and consolidated sleep, along with enhanced starvation tolerance and survival compared to *Drosophila melanogaster*, the developmental trajectory of these phenotypes is unknown. In contrast to adults, *D. mojavensis* larvae sleep less than *D. melanogaster* but exhibit deeper sleep. They also consume more food and survive longer under starvation, linking reduced larval sleep with increased nutrient intake and starvation resistance. Metabolic analyses demonstrate elevated triglyceride accumulation across the *D. mojavensis* lifespan, consistent with enhanced energy storage capacity. Together, these findings reveal a reversal of sleep and feeding phenotypes across development in *D. mojavensis*, raising the possibility that the balance between sleep and other behavioral and metabolic demands is shaped by distinct ecological pressures across life stages.

## Introduction

Sleep is an evolutionarily conserved behavior essential for diverse physiological functions, yet natural sleep duration varies enormously across species ([1]; [2]). These interspecific differences often reflect ecological pressures. Under resource limitations, a fundamental trade-off arises: time spent asleep cannot be spent foraging, making sleep and feeding mutually exclusive behaviors. Many insects and mammals forgo sleep when faced with acute starvation to search for food, underscoring a core conflict between sleep and feeding drives. Consistent with this, satiated animals tend to become quiescent post-feeding, whereas hunger strongly disrupts sleep ([3]). Thus, dynamic regulation of sleep and feeding in response to internal state and environmental conditions is thought to maximize fitness. However, the precise strategies by which animals tune these behaviors, particularly under extreme ecological conditions, remain poorly understood.

Environmental and physiological stressors reshape sleep behavior. Thermal stress inputs, such as extreme heat, activate neural circuits that disrupt sleep by triggering high sleep latency and sleep fragmentation ([4]; [5]). Paradoxically, accumulating evidence suggests that sleep itself can improve survival under such harsh conditions by conserving energy and maintaining homeostasis. Reduced locomotor activity and increased sleep duration have been shown to prolong survival during nutrient deprivation ([6]). A similar pattern has also been observed in the desert-adapted species *Drosophila mojavensis*: these flies maintain high sleep levels during food and water deprivation in contrast to *Drosophila melanogaster* and survive longer under such conditions, supporting the notion that elevated sleep fosters stress resilience and enhances resistance to resource scarcity ([7]). Related, in the nematode *Caenorhabditis elegans*, stress-induced sleep emerges after noxious stress exposure, and sleep-active neurons are essential for mediating sleep’s protective role for survival ([8]; [9]). Taken together, sleep functions as a conserved adaptive response for enhancing stress tolerance.

Nearly all studies of sleep and stress have focused on adulthood. However, developmental sleep is hypothesized to fulfill physiological functions that differ from those in maturity ([10]). Across phylogeny, juveniles consistently sleep more than adults, and this ontogenetic transition in sleep timing is accompanied by functional changes to support stage-specific developmental processes. *Drosophila* larvae exhibit prolonged quiescent periods driven by specific neuromodulatory and neuropeptidergic circuits, facilitating developmental-specific functions such as neurogenesis and neural circuit assembly ([11]; [12]). Similar sleep patterns and functions have been observed in mammals across the lifespan, where neonates spend a greater proportion of time in REM sleep, which gradually declines with age ([13]; [14]). Sleep is thus not a static behavior but a highly plastic state that is restructured throughout development to meet dynamically changing physiological demands. Despite this, it remains unclear how the genetic and neurobiological mechanisms underlying sleep-based resilience in adults are manifested earlier in life. Experimental evolution studies in *D. melanogaster* demonstrate that selection for adult starvation resistance increases adult sleep and reduces adult feeding, while producing developmentally specified changes such as extended development and elevated larval feeding, but little effect on larval sleep, implying stage-specific regulation of these traits ([15]). These stage-specific differences may be particularly relevant for species such as *D. mojavensis*, whose larvae develop within necrotic cactus tissues, while adults encounter the variable resources and environmental stresses characteristic of arid habitats. Here, we use *D. mojavensis* to investigate whether the distinctive sleep phenotype associated with survival in harsh environments reflects a fixed behavioral strategy or is reorganized across development to meet changing ecological demands.

## Results

### Approach for comparing larval sleep across species

We studied *D. melanogaster* (Canton-S; CS) and three geographically distinct *D. mojavensis* populations: Baja, Wrigleyi, and Mojavensis (Moja) (Figure 1A). These species belong to the melanogaster and repleta groups, respectively. In nature, *D. mojavensis* larvae develop within decomposing cactus tissues, where the host plant and associated microorganisms contribute to their nutritional and chemical environment. The populations examined here represent three of the four major host-associated populations: Baja from Baja California, Wrigleyi from Santa Catalina Island, and Moja from the Mojave Desert ([21][22]). Sampling these populations allowed us to examine larval sleep across geographically distinct, host-associated populations within *D.mojavensis*.

**Figure 1.**
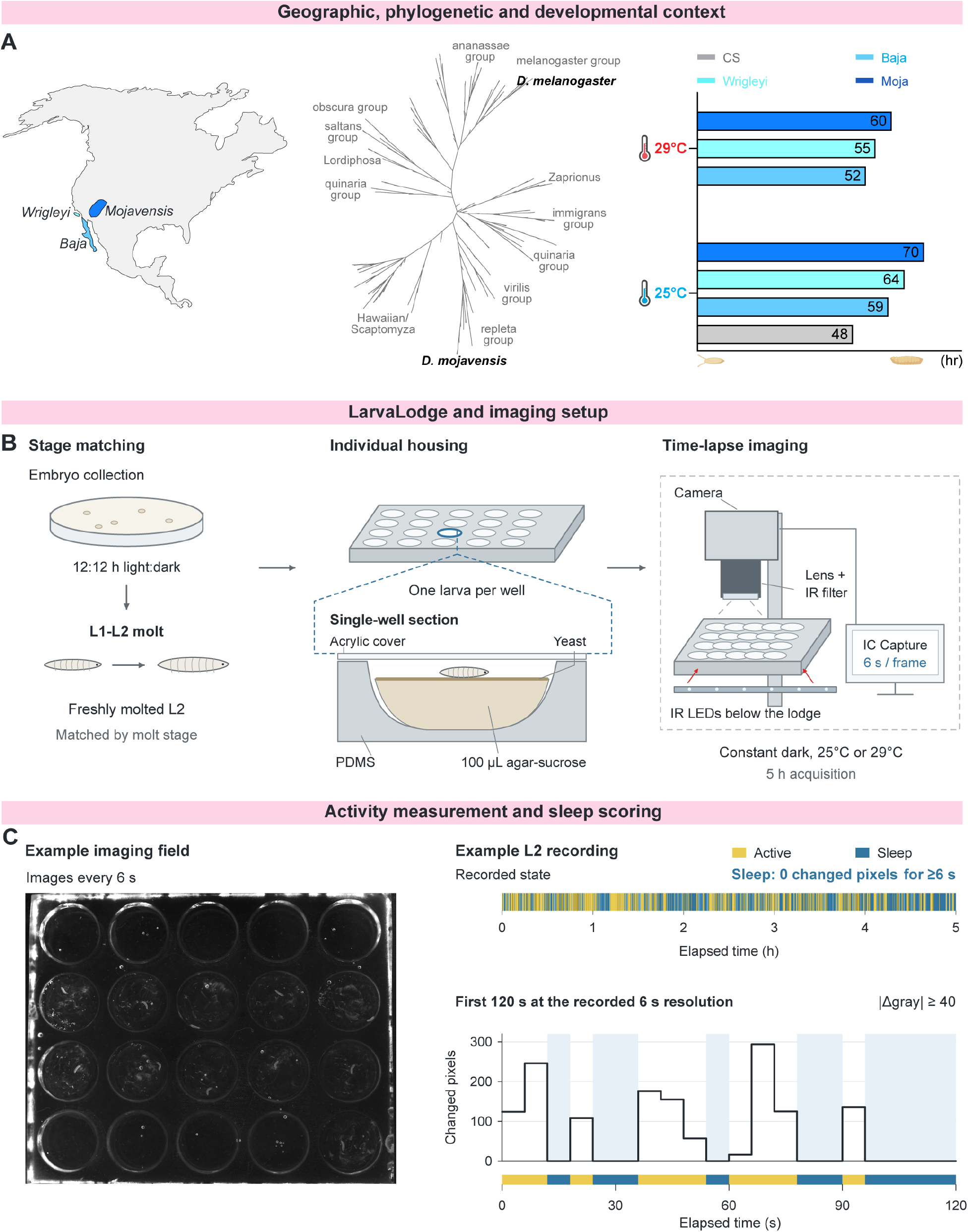
Comparative framework and quantification of larval sleep. **(A)** Left, geographic locations of the three *D. mojavensis* populations examined (Baja, Wrigleyi, and Mojavensis [Moja]). Middle, schematic phylogenetic placement of *D. melanogaster* and *D. mojavensis* within the melanogaster and repleta groups. Right, mean developmental time (hours from embryo to L2) for *D. melanogaster* (*CS*, grey, n=20) and three *D. mojavensis* populations (*Baja*, light blue, n=20; *Wrigleyi*, cyan, n=20; *Moja*, dark blue, n=20) at 25°C and 29°C. **(B)** Stage matching, individual housing, and time-lapse imaging. Embryos were reared under a 12:12 h light:dark cycle, and freshly molted L2 were collected immediately after the L1-L2 molt. Each larva was housed in a separate polydimethylsiloxane (PDMS) well containing 100 µl of 3% agar and 2% sucrose medium topped with yeast paste and covered with a transparent acrylic sheet. Images were acquired through an infrared (IR) filter every 6 s for 5 h in constant darkness at 25°C or 29°C. **(C)** Representative imaging field (left) and example 5-h L2 recording (upper right), with the first 120 s expanded below. Activity was calculated as the number of pixels whose absolute intensity change between consecutive images was ≥40 gray-scale units. Sleep was defined as zero changed pixels for ≥6 s, following the criterion previously validated in *D. melanogaster* larvae. Yellow and blue indicate active and sleep epochs, respectively; pale-blue shading highlights sleep epochs in the expanded activity trace.

To compare larvae at a defined developmental stage, we first measured the time required for embryos to reach the second instar (L2), when we typically assess sleep. At 25°C, mean developmental times were 48 h for CS and 59, 64, and 70 h for Baja, Wrigleyi, and Moja, respectively. Rearing *D. mojavensis* at 29°C shortened these times to 52, 55, and 60 h, bringing developmental timing closer to that of CS at 25°C (Figure 1A). For most experiments, we collected freshly molted L2 immediately after the L1–L2 molt, ensuring that behavioral comparisons were made at the same developmental transition rather than at a fixed chronological age (Figure 1B).

For sleep assays, we used the LarvaLodge approach established for *D. melanogaster* larvae ([11]), which builds on individual-well monitoring and image-analysis methods developed in *C.elegans* ([16]; [17]). Individual larvae were housed in separate wells containing agar-sucrose medium topped with yeast paste and imaged every 6 s for 5 h under infrared illumination in constant darkness at 25°C or 29°C (Figure 1B). Activity was quantified from thresholded changes in pixel intensity between consecutive images, with zero activity indicating no movement detected above the noise threshold (Figure 1C). Our previous work in *D. melanogaster* demonstrated that this imaging approach detects both locomotor and feeding movements, allowing behavioral quiescence to be distinguished from feeding. Periods of larval quiescence lasting ≥6 s meet established behavioral criteria for sleep across phylogeny, including rapid reversibility, increased arousal threshold, characteristic postural changes, and homeostatic rebound following sleep deprivation ([11]). We applied this established definition of sleep to our cross-species comparisons (Figure 1C).

### Sleep is reduced in *Drosophila mojavensis* larvae

Desert-adapted flies exhibit increased sleep in adulthood, which has been associated with enhanced survival under harsh conditions ([7]). However, during earlier developmental larval stages, these animals do not encounter the same degree of food scarcity or temperature stress ([18]; [19]). To investigate whether the distinctive adult sleep phenotype of desert flies is maintained across development, we assessed sleep in *D. mojavensis* L2 under different rearing and assay temperatures. When both species were reared and assayed at 25°C, *D. mojavensis* L2 slept substantially less than *CS* without a corresponding increase in waking activity. *D. mojavensis* L2 sleep was also highly fragmented, characterized by markedly shorter sleep bouts (Figure 2A-D).

**Figure 2.**
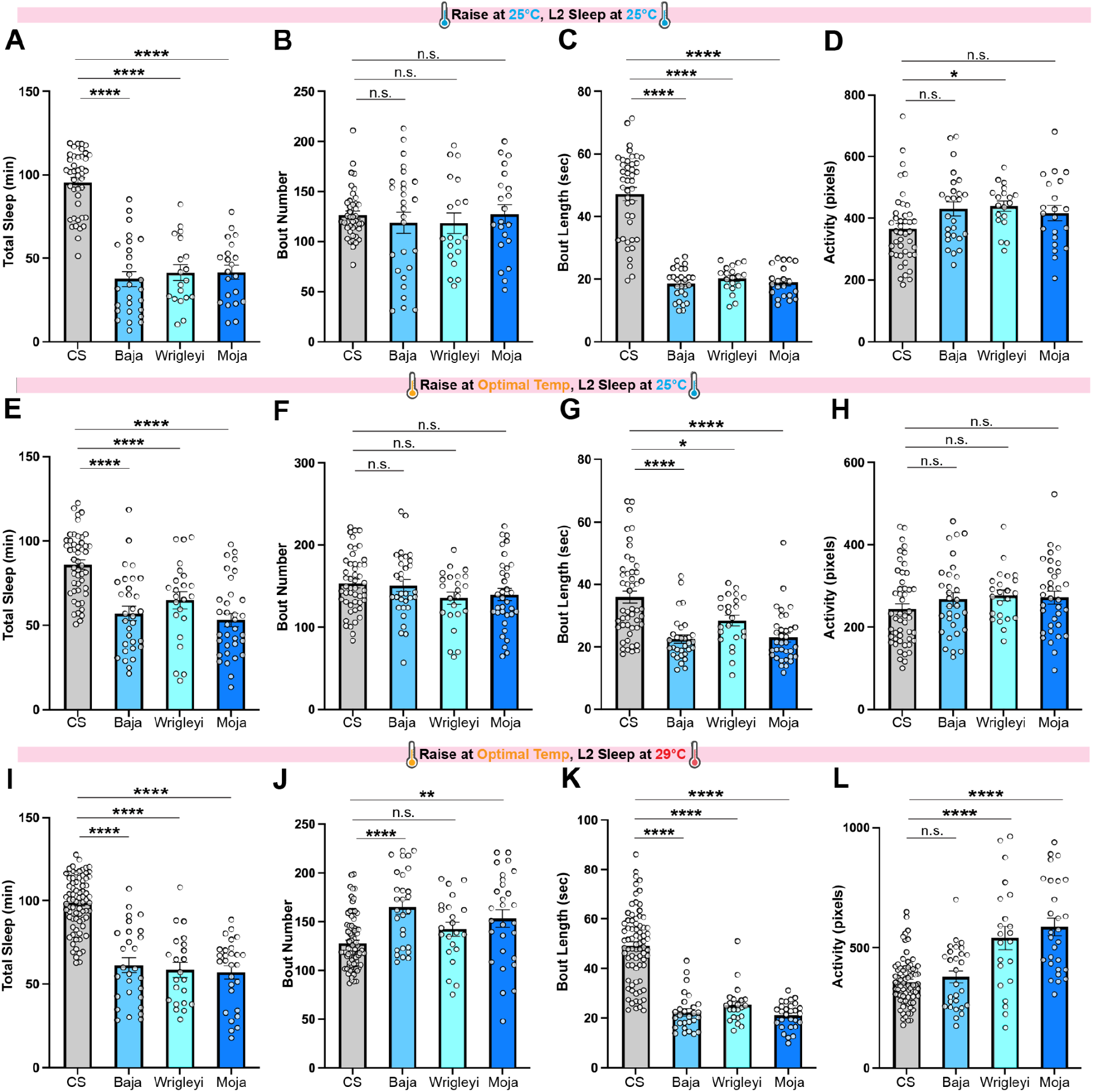
Reduced and fragmented sleep in *D. mojavensis* L2 across temperature conditions. **(A-D)** *D. melanogaster* (*CS,* grey, n=42) and *D. mojavensis* (*Baja*, light blue, n=26; *Wrigleyi*, cyan, n=19; *Moja*, dark blue, n=21) were reared at a uniform 25°C, and L2 sleep was measured over a three-hour interval at 25°C. (A) Total sleep duration. (B) Sleep bout number. (C) Sleep bout length. (D) Waking activity. (A-D) One-way ANOVAs followed by Dunnett’s multiple comparisons test (**** p<0.0001, * p<0.05, n.s. p>0.05). **(E-H)** *D. melanogaster* (*CS,* grey, n=49) and *D. mojavensis* (*Baja*, light blue, n=31; *Wrigleyi*, cyan, n=23; *Moja*, dark blue, n=34) were raised at species-specific temperatures selected to more closely match developmental timing (*D. melanogaster*, 25°C; *D.mojavensis*, 29°C), and L2 sleep was measured over a three-hour interval at 25°C. (E) Total sleep duration. (F) Sleep bout number. (G) Sleep bout length. (H) Waking activity. (E-H) One-way ANOVAs followed by Dunnett’s multiple comparisons test (**** p<0.0001, * p<0.05, n.s. p>0.05). **(I-L)** *D. melanogaster* (*CS,* grey, n=77) and *D. mojavensis* (*Baja*, light blue, n=28; *Wrigleyi*, cyan, n=22; *Moja*, dark blue, n=28) were raised at species-specific temperatures, and L2 sleep was measured over a three-hour interval at 29°C. (I) Total sleep duration. (J) Sleep bout number. (K) Sleep bout length. (L) Waking activity. (I-L) One-way ANOVAs followed by Dunnett’s multiple comparisons test (**** p<0.0001, ** p<0.01, * p<0.05, n.s. p>0.05).

Given the developmental timing differences described above (Figure 1A), we next reared each species at temperatures that more closely aligned their developmental progression (*D. melanogaster*, 25°C; *D. mojavensis*, 29°C) and measured L2 sleep at either 25°C or 29°C. This allowed us to assess whether the reduced sleep phenotype persisted across different rearing and assay temperatures. Under both conditions, *D. mojavensis* L2 consistently showed a reduction in total sleep and shorter sleep bouts than *D. melanogaster*, indicating that the short, fragmented sleep phenotype persists across the temperature conditions tested (Figure 2E-L). Thus, desert-adapted fly strains sleep less during larval stages, in contrast to excess sleep observed in adulthood.

To investigate whether dietary differences might have an impact on sleep, we assessed sleep in L2 raised on an Opuntia cactus-based medium instead of standard agar-yeast food. We quantified sleep architecture in both the first (F1) and fifth (F5) generations after transitioning to the cactus diet to minimize potential short-term effects of transitioning from laboratory food (Figure S1). In both generations, *D. mojavensis* L2 exhibited reduced total sleep and fragmented sleep structure relative to *D. melanogaster*, indicating that the short-sleep phenotype is retained even on the cactus diet. In addition to *D.mojavensis*, comparable sleep phenotypes were also observed in *D. arizonae*, a closely related desert species in the repleta group that shares similar ecological niches. The similarity in sleep traits suggests that reduced larval sleep is not unique to *D. mojavensis* and may extend to closely related desert-adapted species (Figure S2).

### Intact sleep homeostasis and increased sleep depth in *D. mojavensis* larvae

To determine whether *D. mojavensis* L2’s reduced, fragmented sleep reflects impaired sleep homeostasis, we tested sleep rebound following sleep deprivation (SD). A high-intensity blue light pulse was intermittently delivered over a 3 hour window to disrupt sleep in *D. melanogaster* and *D. mojavensis* (Figure 3A). During the subsequent 3-hour recovery period, both sleep-deprived L2 species exhibited rebound sleep compared to non-deprived controls. In *D. melanogaster*, rebound was driven by upward trends in both sleep bout number and bout length; in *D.mojavensis*, rebound was driven by extension in sleep bout length (Figure 3B-C). As expected, waking activity was elevated during deprivation in both species, but during recovery it dipped slightly below baseline in *melanogaster*, consistent with compensatory suppression of activity to facilitate rebound sleep, whereas in *mojavensis* it returned to baseline levels (Figure 3D). These results indicate that despite reduced and fragmented baseline sleep, *D. mojavensis* larvae retain a homeostatic sleep response analogous to that observed in *D. melanogaster* ([11]).

**Figure 3.**
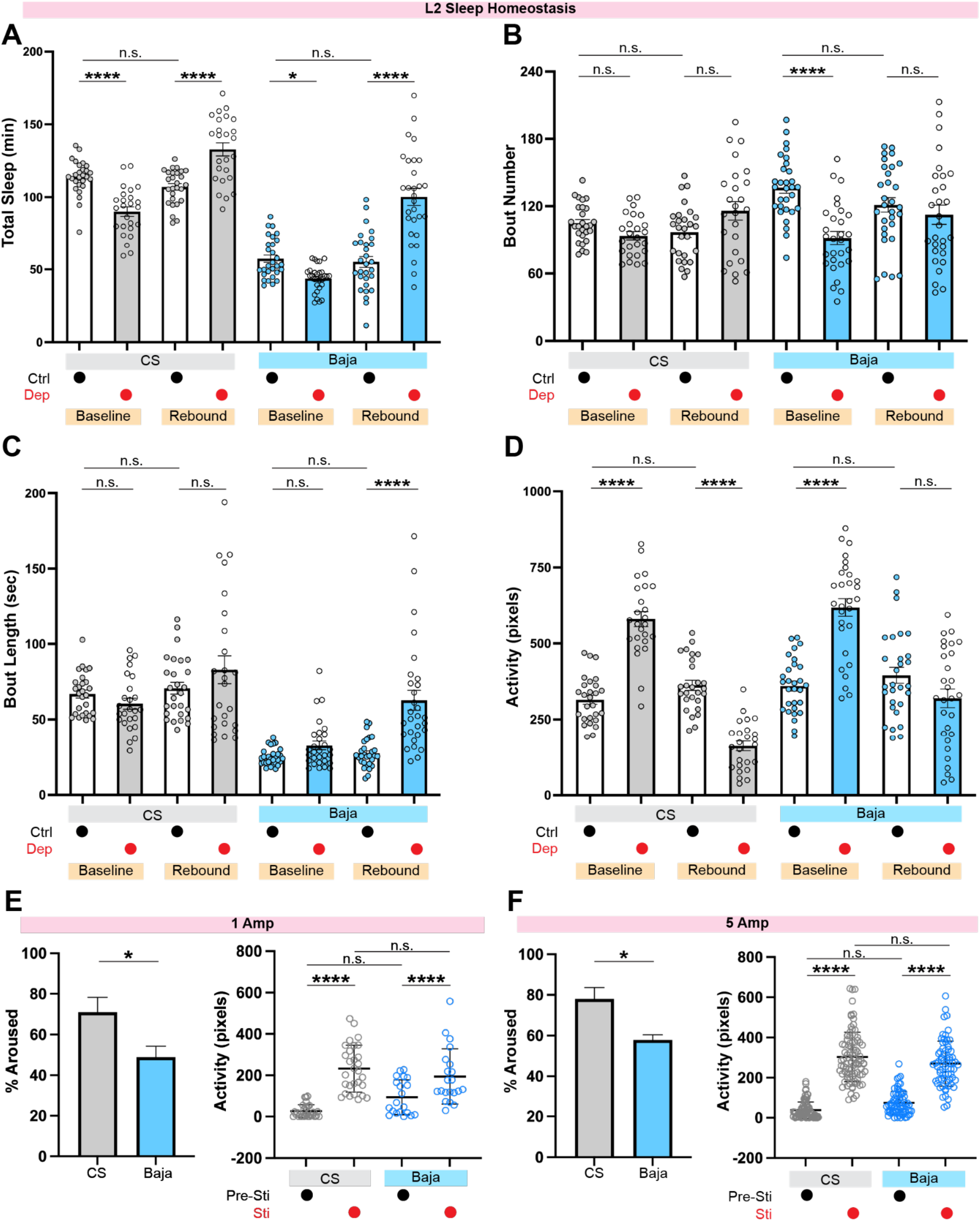
Homeostatic sleep regulation and higher arousal threshold in *D. mojavensis* L2. **(A-D)** *D. melanogaster* (*CS*, grey) and *D. mojavensis* (*Baja*, light blue) L2 were subjected to 3 hours of sleep deprivation (SD) via repetitive light pulses (Baseline) followed by a 3-hour recovery period (Rebound) (*CS* Baseline Ctrl, n=27; *CS* Baseline Dep, n=25; *CS* Rebound Ctrl, n=27; *CS* Rebound Dep, n=25; *Baja* Baseline Ctrl, n=30; *Baja* Baseline Dep, n=29; *Baja* Rebound Ctrl, n=30; *Baja* Rebound Dep, n=29). Quantifications of (A) total sleep duration, (B) sleep bout number, (C) sleep bout length, and (D) waking activity demonstrate intact sleep homeostasis in *D. mojavensis* L2. (A-D) One-way ANOVAs followed by Sidak’s multiple comparisons test (**** p<0.0001, * p<0.05, n.s. p>0.05). **(E)** *D. melanogaster* (*CS*, grey, n=4 replicates) and *D. mojavensis* (*Baja*, light blue, n=5 replicates) L2 were exposed to repetitive low-intensity (1 Amp) light pulses over a 1-hour testing window. The percentage of L2 aroused upon stimulation was measured, and activity levels before (Pre-Sti) and during (Sti) stimulation were recorded to assess locomotor responsiveness to light (*CS* Pre-Sti, n=30; *CS* Sti, n=30; *Baja* Pre-Sti, n=21; *Baja* Sti, n=21). (E) Arousal threshold comparison: unpaired two-tailed Student’s t-test (* p<0.05). Activity quantification: one-way ANOVAs followed by Sidak’s multiple comparisons test (**** p<0.0001, n.s. p>0.05). **(F)** *D. melanogaster* (*CS*, grey, n=3 replicates) and *D. mojavensis* (*Baja*, light blue, n=3 replicates) L2 were exposed to repetitive high-intensity (5 Amp) light pulses over a 1-hour testing window. L2 arousal and activity were quantified as in (E) (*CS* Pre-Sti, n=88; *CS* Sti, n=88; *Baja* Pre-Sti, n=68; *Baja* Sti, n=68). (F) Arousal threshold comparison: unpaired two-tailed Student’s t-test (* p<0.05). Activity quantification: one-way ANOVAs followed by Sidak’s multiple comparisons test (**** p<0.0001, n.s. p>0.05).

We next asked whether reduced sleep quantity in *D. mojavensis* L2 is accompanied by increased sleep depth. We assessed sleep depth via arousal-threshold experiments, delivering 4-second light stimuli of low (1 Amp) and high (5 Amp) intensity every 2 minutes over a 1-hour window. Stimuli were delivered irrespective of behavioral state, and arousal responses were subsequently analyzed only for larvae that were quiescent at the time of stimulus delivery. We observed that *D. mojavensis* L2 were less likely to awaken than *D. melanogaster* at both stimulus intensities (Figure 3E, F), indicating an increased arousal threshold. Baseline activity and locomotor responses to the stimuli were similar between species, arguing against general differences in locomotor responsiveness or light sensitivity as explanations for the reduced arousal response (Figure 3E, F).

### Sleep and feeding exhibit opposing patterns across the lifespan in *D.mojavensis*

Adult *D. mojavensis* exhibit increased sleep together with enhanced starvation resistance, raising the question of how these phenotypes are organized earlier in development when sleep is markedly reduced. We therefore asked whether the reduced larval sleep phenotype is accompanied by altered feeding behavior. Using a dye-based absorbance feeding assay, we first quantified food consumption in L2. When equal numbers of larvae from each species were assessed, *D. mojavensis* groups consumed significantly more food than *D. melanogaster* groups (Figure 4A). To rule out the possibility of a body size confound, we measured weight across the life cycle and found that *D. mojavensis* L2 are approximately half the body weight of *D. melanogaster* at this stage (Figure 4B), indicating that greater food consumption was not due to larger body size. To account for this scaling difference, we repeated the feeding assay under a weight-matched design by doubling the number of *D. mojavensis* L2 so that total biomass matched that of the *D. melanogaster* group. The difference in food consumption was even greater with this design (Figure 4C), confirming increased food intake in *D. mojavensis* L2. We then asked whether these opposing sleep and feeding phenotypes were limited to L2 or persisted later in larval development. In third instar larvae (L3), *D. mojavensis* likewise exhibited reduced and fragmented sleep compared to *D. melanogaster* (Figure 4D-G), along with increased food consumption (Figure 4H).

**Figure 4.**
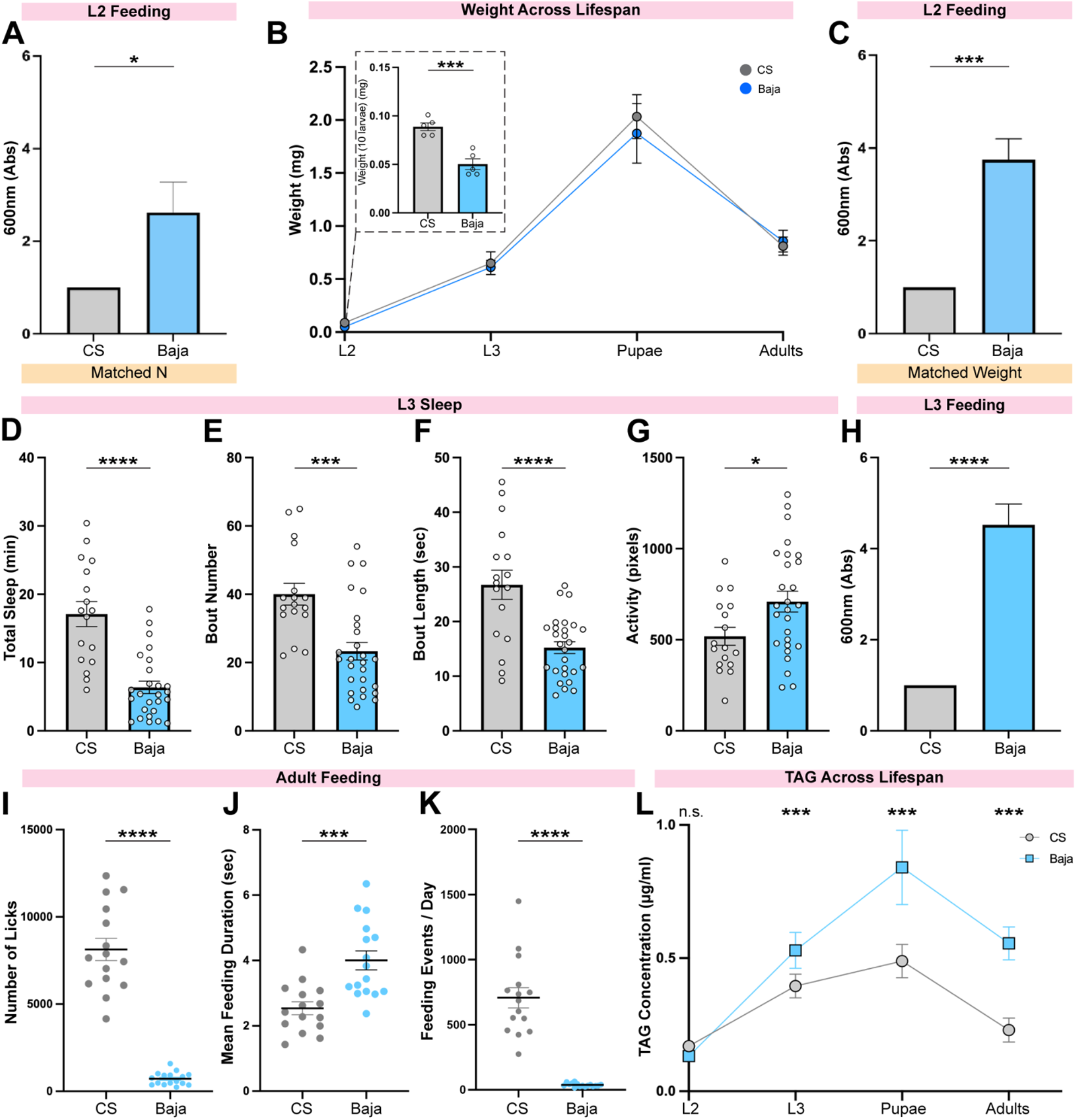
Enhanced larval feeding and developmental energy storage. **(A-C)** *D. melanogaster* (*CS*, grey) and *D. mojavensis* (*Baja*, light blue) L2 were assayed for food consumption using a dye-based absorbance assay. (A) Food consumption under matched sample-size conditions (Matched N; *CS*, 5 independent replicates; *Baja*, 5 independent replicates). Unpaired two-tailed Student’s t-test (* p<0.05). (B) Body weight measurements across life stages (*CS*, n=5 replicates for each stage; *Baja*, n=5 replicates for each stage). Unpaired two-tailed Student’s t-test (*** p<0.001) (C) Food consumption under weight-matched conditions (Matched Weight) (*CS*, n=5 replicates; *Baja*, n=5 replicates). Unpaired two-tailed Student’s t-test (*** p<0.001). **(D-G)** *D. melanogaster* (*CS*, grey, n=17) and *D. mojavensis* (*Baja*, light blue, n=27) L3 were analyzed for sleep. (D) Total sleep duration. (E) Sleep bout number. (F) Sleep bout length. (G) Waking activity. (D-G) Unpaired two-tailed Student’s t-test (**** p<0.0001, *** p<0.001, * p<0.05). **(H)** *D. melanogaster* (*CS*, grey, n=5 replicates) and *D. mojavensis* (*Baja*, light blue, n=5 replicates) L3 were analyzed for food consumption using the dye-based absorbance. Unpaired two-tailed Student’s t-test (**** p<0.0001). **(I-K)** *D. melanogaster* (*CS*, grey) and *D. mojavensis* (*Baja*, light blue) adults were analyzed for feeding interactions using FLIC. (I) The total number of individual food interactions over 24h was significantly elevated in adult *CS* females relative to *Baja*. Welch’s t-test t=11.57, p<0.0001. (J) The mean duration of feeding events that include consecutive licks was prolonged in *Baja* adults compared to *CS*. Welch’s t-test t=4.164, p=0.0003. (K) The total number of feeding events per day was increased in *CS* adult females relative to *Baja*. Welch’s t-test t=8.608, p<0.0001. n=15-16 flies/group for (I-K). **(L)** Whole-body triglyceride (TAG) levels across development (Each biological replicate consisted of 25 L2 larvae, 25 L3 larvae, 5 pupae, or 5 adults; n=6 independent biological replicates per group.). TAG levels were comparable at the L2 stage but were significantly higher in *D. mojavensis* from L3 onward. Unpaired two-tailed Student’s t-test (*** p<0.001).

In stark contrast to larval stages, *D. mojavensis* adults exhibit elevated and consolidated sleep compared to *D. melanogaster*. We thus investigated feeding behavior at this stage using a liquid food interaction assay and found that adult *D. mojavensis* exhibited fewer individual food interactions and feeding events than *D. melanogaster*, although the mean duration of individual feeding events was longer (Figure 4I-K) ([20]). Because larval and adult feeding were assessed using different, stage-appropriate assays, these measurements establish relative species differences within each developmental stage rather than directly comparable measures of food intake across stages. To determine whether these behavioral differences are accompanied by differences in energy storage, we measured triglyceride (TAG) levels across development. *D. mojavensis* accumulated more lipid reserves than *D. melanogaster* beginning at the L3 stage, a difference that persisted into adulthood (Figure 4L). Thus, increased feeding is evident before the species difference in lipid stores emerges, although the relationship between these behavioral and metabolic phenotypes remains to be determined.

### Differential starvation responses and enhanced survival in *D. mojavensis* L2

We next examined behavioral responses to acute food deprivation in *D. mojavensis* L2. In a food-deprived starvation paradigm, both *D. mojavensis* and *D. melanogaster* L2 showed the expected response of suppressing sleep relative to fed controls, consistent with hunger-induced foraging behavior. However, closer examination revealed that *D. mojavensis* L2 suppressed sleep to a lesser extent than *D. melanogaster* under starvation, including greater sleep duration, more sleep bouts, and longer bout length (Figure 5A-C). Thus, while both species suppress sleep in the absence of food, *D. mojavensis* L2 do not curtail sleep as drastically as *D. melanogaster*.

**Figure 5.**
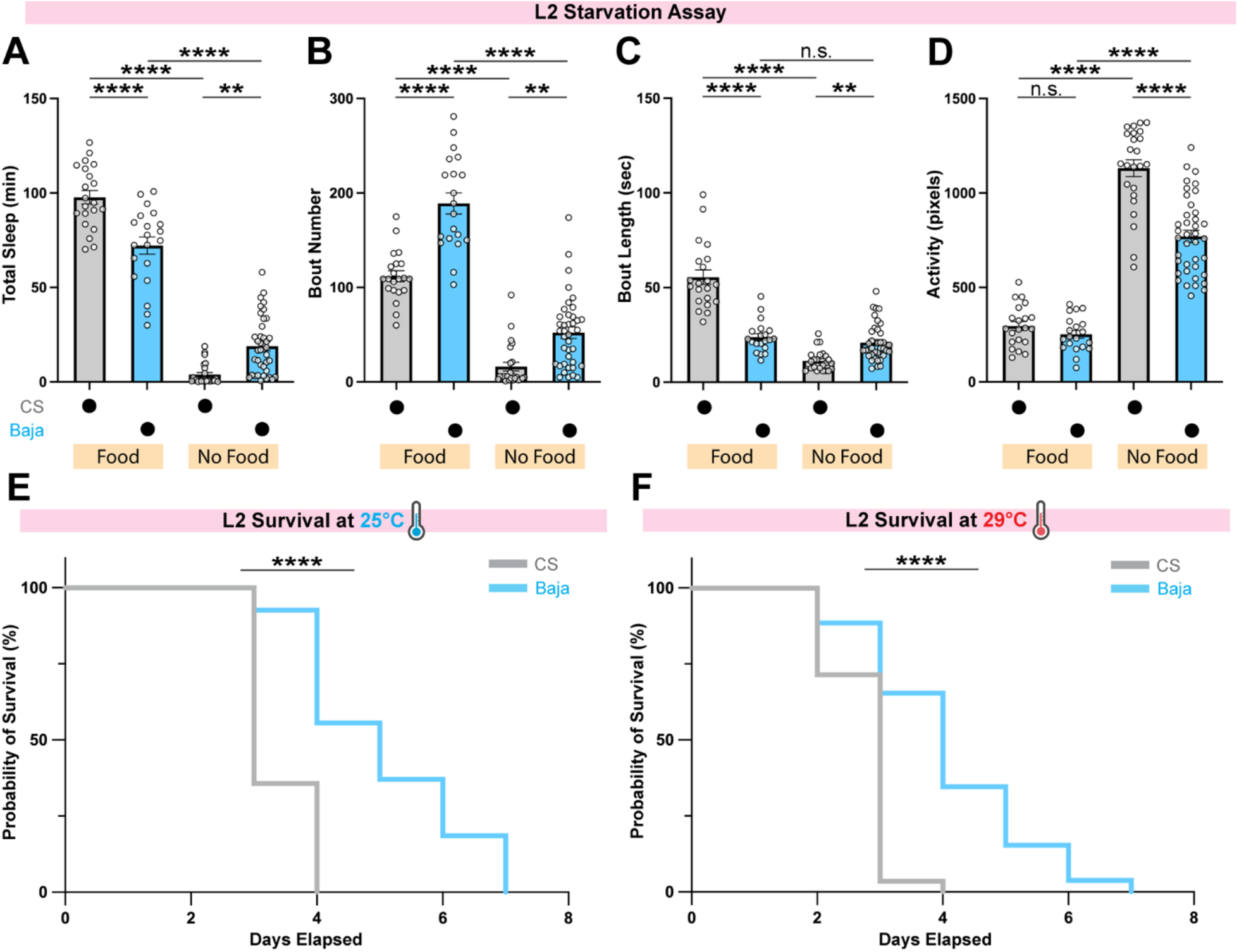
*D. mojavensis* L2 exhibit differential behavioral responses to starvation and enhanced survival. **(A-D)** *D. melanogaster* (*CS*, grey, n=21 for “Food”, and n=24 for “No Food”) and *D. mojavensis* (*Baja*, light blue, n=20 for “Food”, and n=39 for “No Food”) L2 were analyzed for sleep under fed and starvation conditions. (A) Total sleep duration. (B) Sleep bout number. (C) Sleep bout length. (D) Waking activity. (A-D) One-way ANOVAs followed by Tukey’s multiple comparisons test (**** p<0.0001, *** p<0.001, ** p<0.01). **(E-F)** Survival of *D. melanogaster* (*CS*, grey) and *D. mojavensis* (*Baja*, light blue) L2 larvae under starvation conditions. (E) Survival at 25°C. *D. mojavensis* L2 (*Baja*, n=27) survived significantly longer than *D. melanogaster* L2 (*CS*, n=28) under starvation. (F) Probability of survival at 29°C. *D. mojavensis* L2 (*Baja*, n=26) survived significantly longer than *D. melanogaster* L2 (*CS*, n=28) under starvation. (E-F) Kaplan–Meier survival analysis with log-rank (Mantel–Cox) tests (**** p<0.0001). Experiments were performed in 2 independent replicates, with a total of n=26-28 larvae per species at each temperature.

Is the enhanced starvation resistance previously observed in adult *D. mojavensis* also present during larval development? Under starvation conditions, *D. mojavensis* L2 also survived longer than *D. melanogaster* L2 at both 25°C (Figure 5E) and 29°C (Figure 5F), demonstrating that enhanced starvation resistance is already evident during larval development. Notably, the behavioral response to starvation differs across developmental stages: while *D. mojavensis* larvae suppress sleep in the absence of food, adults maintain elevated sleep rather than exhibiting the starvation-induced sleep suppression observed in adult *D. melanogaster*. Thus, enhanced starvation resistance in *D. mojavensis* is present at both larval and adult stages despite distinct stage-specific sleep responses to food deprivation.

## Discussion

Here we report a developmental reversal of sleep and feeding phenotypes in the desert-adapted *D. mojavensis*. Larvae exhibit reduced and fragmented sleep alongside markedly elevated feeding, whereas adults exhibit increased and consolidated sleep together with less frequent feeding. Despite these opposing behavioral phenotypes, *D. mojavensis* larvae, like adults, exhibit enhanced starvation resistance relative to *D. melanogaster*. Thus, the relationship among sleep, feeding, and starvation resistance differs substantially across life stages within the same species.

*D. mojavensis* is a cactophilic species that feeds and breeds in necrotic cactus tissues, and distinct geographic populations use different cactus hosts across the species range. These substrates are dynamic food sources; cactus necroses are chemically and temporally volatile with microbial environments that change across fermentation stages. Thus, larvae develop and feed within decaying cactus tissues whose nutritional and chemical composition varies over time, whereas adults must search for, assess, and move among cactus resources while tolerating exposed desert conditions, including heat and desiccation stress ([21–23]). These distinct life-stage ecologies raise the possibility that the opposing sleep and feeding phenotypes we observe reflect different behavioral and energetic demands across development. Reduced larval sleep could increase opportunities for feeding during a period of rapid growth, whereas elevated adult sleep could reduce energetic and water expenditure under conditions in which resources are limited and/or unstable.

Our experiments do not establish that reduced larval sleep causes increased feeding, lipid storage, or starvation resistance. Rather, these phenotypes co-occur in *D. mojavensis*, providing an avenue for testing whether they are functionally linked. Reduced sleep could facilitate increased feeding opportunity, but sleep and feeding could also be independently regulated components of a broader developmental program, and metabolic differences could themselves influence each behavior. Differences in developmental timing between species could also contribute to the observed feeding and metabolic phenotypes, independent of differences in sleep. Direct manipulation of sleep across species will help distinguish among these possibilities.

Reduced and fragmented larval sleep was also observed in *D. arizonae*, indicating that this phenotype is not unique to *D. mojavensis*. Together with previous work demonstrating elevated adult sleep in *D.* arizonae and other desert-associated *Drosophila*, this observation suggests that contrasting larval and adult sleep phenotypes extend beyond *D. mojavensis* ([7]). However, broader phylogenetic sampling and matched larval-adult comparisons will be required to determine whether this pattern reflects shared ancestry, ecological convergence, or species-specific variation.

The persistence of elevated arousal threshold across life stages suggests that deeper sleep may be a lifelong feature of *D. mojavensis* sleep despite opposing larval and adult sleep-quantity phenotypes. These opposing sleep duration phenotypes across life stages argue against a constitutive change in sleep regulation as the basis for elevated adult sleep in *D. mojavensis*. If elevated adult sleep resulted simply from changes in broadly acting sleep-regulatory pathways, larval sleep would be similarly increased. Instead, our findings point toward developmental regulation of the species-specific sleep phenotype. This interpretation is consistent with previous work showing that selection for adult sleep or starvation resistance in *D. melanogaster* does not produce parallel changes in larval behavior ([6], [15]). Neurobiological evidence provides a potential basis for such stage-specific regulation: in *D. melanogaster*, the neuromodulatory mechanisms governing sleep differ across development ([11]). Developmentally distinct sleep-regulatory mechanisms could allow evolutionary changes in sleep to be expressed differently across life stages. Such developmental flexibility could provide a basis for the opposing larval and adult sleep phenotypes observed in *D. mojavensis*.

More broadly, our work contributes to a growing body of evidence that sleep is a flexible, ecologically shaped behavior. Across taxa, animals adjust sleep timing and duration to accommodate life-history events such as migration, breeding, and seasonal foraging. White-crowned sparrows reduce sleep dramatically during migration while maintaining cognitive performance, and Arctic sandpipers virtually eliminate sleep during mating season to maximize reproductive success ([24];[25];[26]). In insects, male *Drosophila* suppress nighttime sleep to court females ([27];[28]). These examples show that animals can sacrifice sleep when ecological demands take priority. *D. mojavensis* larvae may represent a developmental parallel, in which reduced sleep occurs alongside increased feeding during a period of rapid growth. An important next step will be to identify the neurogenetic mechanisms underlying these stage-specific differences in sleep and feeding. Given that neuromodulators such as dopamine, octopamine, serotonin, and NPF are known to regulate sleep and feeding in adults, one possibility is that developmental differences in these systems contribute to the opposing sleep and feeding phenotypes observed across the *D. mojavensis* lifespan ([29];[30]). Taken together, our findings reveal a striking developmental reversal in sleep and feeding phenotypes in *D. mojavensis*. This work supports the view that sleep is not static but a highly plastic state that can be reorganized across life stages in relation to changing environmental and physiological demands.

## Methods and Materials

### Adult Rearing

Fly stocks were cultured on standard cornmeal molasses media (per 1L H2O: 12 g agar, 29 g Red Star yeast, 71 g cornmeal, 92 g molasses, 16mL methyl paraben 10% in EtOH, 10mL propionic acid 50% in H2O) at 25°C with 60% relative humidity and entrained to a daily 12h light, 12h dark schedule. *Canton-S* (*CS*) were provided by Dr. Gero Miesenböck (University of Oxford). Primary stocks *D.mojavensis*.*baja* (collected March 2020, La Paz, Mexico), *D. mojavensis.moja* (collected February 2020, North Joshua Tree National Park, CA), *D.mojavensis.wrigleyi* (collected November 2017, Catalina Island, CA) were a gift from Dr. Luciano Matzkin (University of Arizona). *D. arizonae* were ordered from the National *Drosophila* Species Stock Center (SKU: 15081-1271.36).

### Larval Rearing

All fly stocks were cultured on a standard molasses medium and raised in incubators on a 12:12 light:dark (LD) cycle. Different rearing temperatures were used across species to synchronize their developmental time: *CS* was maintained at 25℃; *baja*, *moja*, *wrigleyi*, and *arizonae* at 29℃. To collect synchronized L2, adult flies were placed in an embryo collection cage (Genesee Scientific, cat#: 59-100) and eggs were laid on a petri dish containing 3% agar, 2% sucrose, and 2.5% apple juice with yeast paste on top.

### Larval Body Weight Measurements

For weight, groups of 10 L2, 5 L3, pupae, or adults were washed in Milli-Q water and dried using a Kimwipe. The samples were then weighed as a group on a scale, and the weight in milligrams (mg) was recorded.

### Larval Sleep Assays and Starvation Assays

On the day of the sleep assay, molting first instar larvae (L1) were collected and moved to a separate dish to complete the molt to L2. Freshly molted L2 larvae were placed into individual wells of the LarvaLodge containing 100 µl of 3% agar and 2% sucrose medium covered with a thin layer of yeast paste. Experiments with Banana-Opuntia medium used a recipe from the National *Drosophila* Species Stock Center (NDSSC; Cornell University): per 1L H2O, 14.16g agar, 27.5 g yeast, 2.23g methyl paraben, 137.5g blended bananas, 95g Karo Syrup, 30g Liquid Malt Extract, 22.33g 100% EtOH, 2.125g powdered opuntia cactus.

For starvation assays, 100 µl of 3% agar-only medium was applied to each well. The LarvaLodge was covered with a transparent acrylic sheet brushed with a thin layer of detergent to prevent condensation, and moved to a DigiTherm incubator at either 25°C or 29°C for imaging in constant dark.

### LarvaLodge Image Acquisition and Processing

Images were captured every 6 sec for 5 hrs with an Imaging Source DMK 23GP031 camera equipped with a Fujinon lens with a Hoya 49mm R72 Infrared Filter. We used IC Capture (The Imaging Source) to acquire time-lapse images. All experiments were carried out in the dark using infrared LED strips positioned below the LarvaLodge.

Images were analyzed using custom-written MATLAB software. Temporally adjacent images were subtracted to generate maps of pixel value intensity change. A binary threshold was set such that individual pixel intensity changes that fell below 40 gray-scale units within each well were set equal to zero (“no change”) to eliminate noise. Pixel changes greater than or equal to the threshold value were set equal to one (“change”). Activity was then calculated by taking the sum of all pixels changed between images. Sleep was defined as behavioral quiescence lasting ≥6 sec, an operational definition previously validated in *Drosophila* larvae based on increased arousal threshold and homeostatic rebound ([11]). Waking activity was calculated from activity values during waking epochs only, excluding periods classified as sleep.

### Arousal Threshold, Light Responsiveness, and Sleep Deprivation via Blue Light Stimulation

Blue light stimulation was delivered using two high-power LEDs (Luminus Phatlight PT-121, 460 nm peak wavelength, Sunnyvale, CA) secured to an aluminum heat sink. The blue light stimulus was delivered for 4 sec every 2 min for 1 hr beginning the 2^nd^ hr after the molt to L2. Stimuli were delivered to all larvae irrespective of behavioral state. Arousal responses were subsequently analyzed only for larvae that were quiescent during the 6-sec interval immediately preceding stimulus delivery. A larva was scored as aroused if locomotor activity was detected during the 6-sec interval immediately following stimulation, and the percentage of quiescent larvae that aroused in response to each stimulus was calculated. Low-and high-intensity stimuli were delivered at 1 Amp and 5 Amp, respectively. To assess whether differences in arousal reflected general differences in locomotor responsiveness to blue light, activity was quantified during the 30 sec before and 30 sec after stimulation.

For sleep deprivation, we used a 5-Amp stimulus for 18 sec every 30 sec for 3 hrs beginning 1 hr after the molt to L2. Undisturbed control larvae were placed in a separate incubator under identical conditions without blue light exposure during the deprivation.

### Feeding Behavior Analysis

Freshly molted L2 were placed in a petri dish containing blue-dyed 3% agar, 2% sucrose, and 2.5% apple juice with blue-dyed yeast paste on top for 4 hrs at 25°C in constant darkness. After 4 hrs, L2 were washed in water, transferred to microtubes, and frozen at -80°C overnight. Samples were homogenized in 300 µl of distilled water and centrifuged for 5 min at 13,000 rpm. The amount of blue dye in the supernatant was then measured using a spectrophotometer (OD629). OD629 was used as a measure of relative food consumption. This assay measures cumulative food consumption over the 4-hr feeding period and does not distinguish feeding rate during wakefulness from differences in the amount of time available for feeding.

### Triglyceride (TAG) and Bicinchoninic Acid Protein (BCA) Assays

Triglyceride levels were measured using the Stanbio Triglyceride LiquiColor Kit (EMSCO/Fisher, 2100-430) from *Drosophila* adult flies, pupae, and larvae. For each biological replicate (n=6), samples consisted of 25 larvae, 5 pupae, or 5 adults. Samples were collected and washed in cold phosphate-buffered saline (PBS) and homogenized 3 x 10 sec each in 125 µL of lysis buffer (140 mM NaCl, 50 mM Tris-HCl [pH 7.4], 0.04% Triton X-100, and 1X protease inhibitors). The homogenates were centrifuged at 13,000 rpm for 10 min at 4°C, and the supernatants were stored at -20°C until analysis. TAG levels (µg/µL) were determined according to the kit protocol and measured using a microplate reader at 500 nm. TAG levels were normalized to total protein concentration, measured at 562 nm using the Pierce BCA Protein Assay Kit (Cell Center/Thermo, PI23227).

### Survival with Starvation Analysis

Freshly molted second-instar (L2) larvae were placed individually into LarvaLodge wells containing 100 μL of 3% agar and maintained in a DigiTherm incubator at either 25°C or 29°C under constant darkness. Images were acquired at 5-min intervals until all larvae in the LarvaLodge had died. Death was defined as complete and sustained cessation of detectable movement, confirmed by manual inspection of successive images. Every 24 hours, quantitative activity data were cross-checked against the corresponding images for each larva to verify survival scoring. The transparent acrylic lid covering the wells was replaced every 24 hours to minimize condensation and associated artifacts in automated movement detection. When a larva migrated into an adjacent occupied well, data from both larvae were excluded from survival analysis because their movements could no longer be reliably attributed to individual animals. Data were also excluded for larvae in wells where condensation prevented reliable movement detection and survival scoring. Experiments were performed in 2 independent replicates, with a total of n = 26-28 larvae per species and temperature condition.

### FLIC Feeding Assay

Fly Liquid Interaction Counter (FLIC; Sable Systems, North Las Vegas, NV) feeding assay was performed to obtain data on feeding duration and frequency previously described ([20]). 4-8 day old female flies were individually loaded into separate chambers of the *Drosophila* Feeding Monitors (DFM) to track feeding behavior. DFM feeding wells were loaded with a solution of 5.0% sucrose and 50 mg/L MgCl2 in distilled water. Individual flies are aspirated into separate DFM chambers between ZT0-2 and behavior data is recorded for 24h. Feeding data includes measurements from two independent experiments. The DFM detects feeding signals from individual flies via a microcontroller circuit board on each well. Raw feeding signal data is coordinated and processed in the Master Control Unit (MCU). Feeding data was analyzed in R Studio using previously described analysis scripts (https://github.com/PletcherLab/FLIC_R_Code) ([31]).

### Statistical Analysis

All statistical analysis was done in GraphPad (Prism). For comparisons between two conditions, two-tailed unpaired *t*-tests were used. Welch’s *t*-tests were used when unequal variances were assumed, as indicated in the figure legends. For comparisons between multiple groups, ordinary one-way ANOVAs followed by Dunnett’s, Tukey’s, or Sidak’s multiple comparisons tests were used, as indicated in the figure legends. Survival curves were analyzed using Kaplan–Meier survival analysis with log-rank (Mantel–Cox) tests. Statistical tests and sample sizes for individual experiments are indicated in the corresponding figure legends.

## Acknowledgements

We thank members of the Kayser Lab for helpful discussions and input.

## Author contributions

Conceptualization: All authors

Investigation: SL, MS, CN, SHT

Writing – Original Draft: SL, MSK

Writing – Review and Editing: All authors

Project Supervision and Funding: MSK

## Data and materials availability

All data needed to evaluate the conclusions in the paper are present in the paper and/or the Supplementary Materials.

---

**Figure S1.**
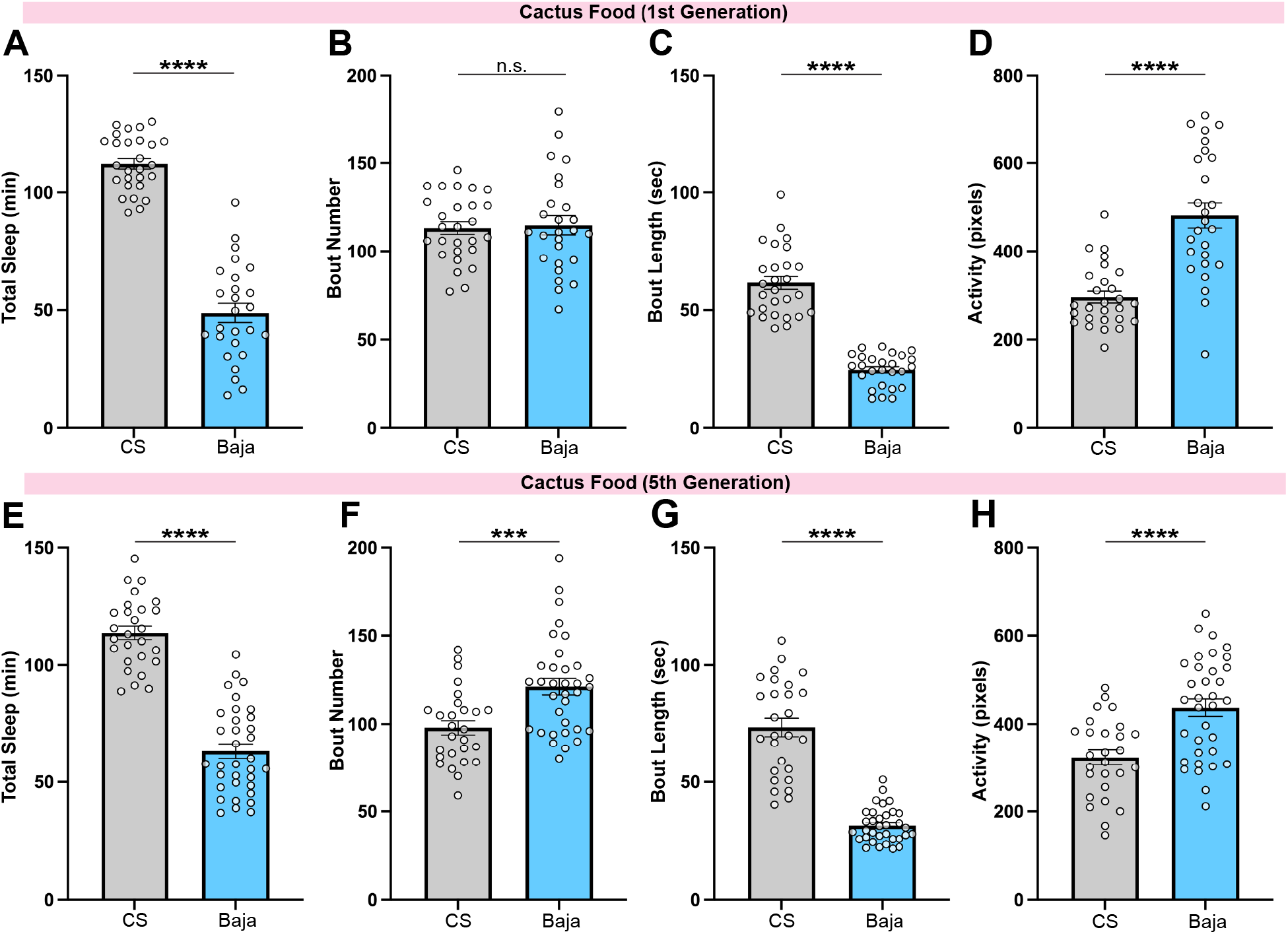
Reduced and fragmented sleep in *D. mojavensis* L2 persists on cactus-based food across generations. **(A-D)** *D. melanogaster* (*CS,* grey, n=27) and *D. mojavensis* (*Baja*, light blue, n=26) P0 generation were raised on banana-Opuntia cactus medium. L2 sleep was measured over a 3-hour interval in the F1 generation. (A) Total sleep duration. (B) Sleep bout number. (C) Sleep bout length. (D) Waking activity. (A-D) Unpaired two-tailed Student’s t-test (**** p<0.0001, n.s. p>0.05). **(E-H)** *D. melanogaster* (*CS,* grey, n=27) and *D. mojavensis* (*Baja*, light blue, n=34) P0 generation were raised on banana-Opuntia cactus medium. L2 sleep was measured over a 3-hour interval in the F5 generation (after five generations on the cactus diet) to assess whether the sleep phenotype persisted across multiple generations on this diet. (E) Total sleep duration. (F) Sleep bout number. (G) Sleep bout length. (H) Waking activity. (E-H) Unpaired two-tailed Student’s t-test (**** p<0.0001, *** p <0.001).

**Figure S2.**
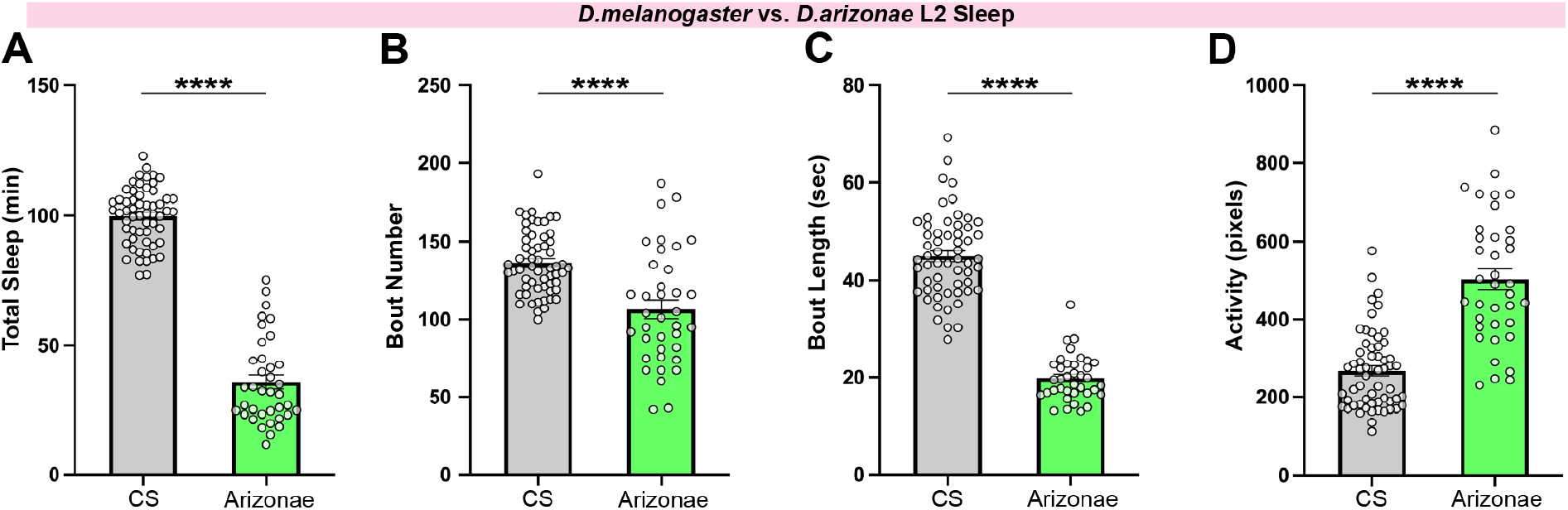
Reduced and fragmented L2 sleep in the closely related desert species D. arizonae. **(A-D)** *D. melanogaster* (*CS,* grey, n=58) and *D. arizonae* (green, n=37) L2 larvae were monitored for 3 hours to assess sleep. (A) Total sleep duration. (B) Sleep bout number. (C) Sleep bout length. (D) Waking activity. (A-D) Unpaired two-tailed Student’s t-test (**** p<0.0001).

